# Early life challenge enhances cortisol regulation in zebrafish larvae

**DOI:** 10.1101/2024.08.11.607500

**Authors:** Luis A. Castillo-Ramírez, Ulrich Herget, Soojin Ryu, Rodrigo J. De Marco

## Abstract

The hypothalamic-pituitary-adrenal (HPA) axis in mammals and the hypothalamic-pituitary-interrenal (HPI) axis in fish are open systems that adapt to the environment during development. Little is known about how this adaptation begins and regulates early stress responses. We used larval zebrafish to examine the impact of prolonged forced swimming at 5 days post-fertilization (dpf), termed early-life challenge (ELC), on cortisol responses, neuropeptide expression in the nucleus preopticus (NPO), and gene transcript levels. At 6 dpf, ELC-exposed larvae showed normal baseline cortisol but reduced reactivity to an initial stressor. Conversely, they showed increased reactivity to a second stressor within the 30-minute refractory period, when cortisol responses are typically suppressed. ELC larvae had fewer corticotropin-releasing hormone (*crh*), arginine vasopressin (*avp*), and oxytocin (*oxt*)-positive cells in the NPO, with reduced crh and avp co-expression. Gene expression analysis revealed upregulation of genes related to cortisol metabolism (*hsd11b2*, *cyp11c1*), steroidogenesis (*star*), and stress modulation (*crh*, *avp*, *oxt*). These results suggest that early environmental challenge initiates adaptive plasticity in the HPI axis, tuning cortisol regulation to balance responsiveness and protection during repeated stress. Future studies should explore the broader physiological effects of prolonged forced swimming and its long-term impact on cortisol regulation and stress-related circuits.

**Summary:** This study explores how early-life challenges in zebrafish affect stress responses and hormone regulation, offering insights into developmental adaptability and stress management mechanisms.

## INTRODUCTION

The HPA/I axis functions as an adaptive system closely linked to bodily responses to environmental challenges that disrupt homeostasis and trigger protective adaptations (Wendelaar Bonga, 1997; Charmandari et al., 2005; Chrousos, 2009). As an open system, it continually adjusts to interactions between the organism and its environment throughout development, enhancing the ability to cope with stress. However, excessive or poorly timed stress can lead to high allostatic load, accelerating maladaptive processes (de Kloet et al., 2005; Danese and McEwen, 2012; Chen and Baram, 2016). While the effects of stress in adulthood are often transient, early-life stress can alter brain development, resulting in long-lasting behavioural changes (Danese and McEwen, 2012; Bick and Nelson, 2016; Teicher et al., 2016; Agorastos et al., 2019). These behavioural symptoms may manifest in childhood, adolescence, or later in adulthood, when the delayed effects of stress-induced alterations in brain development become apparent (Lupien et al., 2009). Additionally, human infants exposed to high cortisol levels *in utero* show elevated baseline cortisol and reduced cortisol responses to separation stress (O’Connor et al., 2013). Similarly, rodent studies show that early postnatal dexamethasone treatment reduces corticosterone responses to stress later in life (Felszeghy et al., 2000), while prolonged exposure to low-dose corticosterone further suppresses reactivity (Kinlein et al., 2015). These findings illustrate how both pre- and post-natal environments shape stress regulation and long-term HPA axis function, although the exact mechanisms remain unclear.

Larval zebrafish (*Danio rerio*) are a valuable model for exploring HPA/I axis adaptation during early development due to their genetic accessibility and external development, which facilitate the analysis of early-life stress (ELS) (Alsop and Vijayan, 2008; Alsop and Vijayan, 2009; Egan et al., 2009; Champagne and Richardson, 2013; Biran et al., 2015; Eachus et al., 2021; Tan et al., 2022). As a non-mammalian vertebrate model (Grunwald and Eisen, 2002; Holtzman et al., 2016), zebrafish have become prominent in ELS research, offering practical advantages such as small size, high reproductive capacity, and low maintenance costs. Their external development and lack of parental care allow for controlled experiments without the confounding effects of prenatal stress. The first stage of stress processing in zebrafish occurs in the nucleus preopticus (NPO), which is homologous to the mammalian paraventricular nucleus (PVN) (Herget et al., 2014). The process involves key neuropeptides like corticotropin-releasing hormone (CRH), which stimulates adrenocorticotropic hormone (ACTH) production in the pituitary, leading to glucocorticoid (GC) release from the interrenal gland, functionally similar to the adrenal gland in mammals (Alderman and Bernier, 2009). The increased secretion of GCs like cortisol after the onset of stress, known as glucocorticoid reactivity (GC_R_), is crucial for the organism’s response to challenge. Beyond their role in stress, GCs regulate a wide range of bodily functions, including metabolism, maintenance of water and electrolyte balance, immunity, growth, cardiovascular function, mood, cognition, reproduction, and development. Primarily produced in the adrenal cortex, along with aldosterone and dehydro-epi-androsterone, all derived from cholesterol, GCs are also synthesized at extra-adrenal sites such as the thymus, brain, and epithelial barriers. These localized productions contribute to spatial specificity in steroid actions, operating independently of systemic and stress-induced fluctuations (Munck et al., 1984; Chrousos, 1998; de Kloet et al., 1998; McEwen, 2007; Timmermans et al., 2019; de Kloet and Joëls, 2023). Together, these features make zebrafish a powerful handle for investigating the molecular mechanisms by which early environments influence brain development and behaviour, as well as for exploring potential therapeutic interventions for stress-related disorders (Veenstra-VanderWeele and Warren, 2015; Bick and Nelson, 2016).

By 4-6 dpf, when the HPI axis becomes functional (Alsop and Vijayan, 2008; Alderman and Bernier, 2009; De Marco et al., 2016), zebrafish larvae in laboratory settings primarily encounter visual cues from conspecifics and mechanosensory inputs from other larvae, unless raised in isolation, along with self-generated motion. Temperature, illumination, and food availability can be tightly controlled, creating an environment with limited variability, ideal for analyzing early HPI axis calibration to external stimuli. For example, their response to water currents, where they adjust swimming to match flow strength, allows for environments with varying physical challenges. Zebrafish larvae adapt to water vortex flows through rheotaxis, orienting themselves against the current, which imposes high energy demands and activates the HPI axis. Controlled vortex flows at specific revolutions per minute (rpm) induce a rapid and significant rise in cortisol levels, unlike conditions without flow, establishing vortex flow as an effective stressor (Castillo-Ramírez et al., 2019). While the metabolic or cardiorespiratory effects of vortex exposure have not yet been measured, the cortisol increase indicates significant energy demands and physiological stress. Additionally, vortex flows offer several advantages as a stressor, minimizing confounding variables such as salt concentration or pH changes and utilizing an innate, reproducible behaviour, reducing experimental variability. Consequently, vortices and less predictable water motions have been used to study stress responses in zebrafish larvae (De Marco et al., 2014; vom Berg-Maurer et al., 2016; Ryu and De Marco, 2017; Castillo-Ramírez et al., 2019; Langebeck-Jensen et al., 2019; Herget et al., 2023; Castillo-Ramírez et al., 2024). Notably, as vortex strength (rpm) increases, cortisol levels rise proportionally, as does larval swimming effort to counteract the flow (Castillo-Ramírez et al., 2019). This quantifiable relationship between vortex intensity and GC_R_ allows precise categorization of stress levels, making vortex flows ideal for repeated stress assays. Using these highly controlled water vortices, we established a high-throughput forced swim test for larval zebrafish. This test revealed that early-life challenge (ELC), in the form of sustained involuntary swimming, triggers prolonged HPI axis activation and alters the cortisol response to a single instance of either homotypic or heterotypic stress 1-4 days post-ELC. ELC also enhances spontaneous activity, reduces startle reactivity, and improves energy efficiency during rheotaxis (Castillo-Ramírez et al., 2019). The mechanisms underlying these effects remain unknown.

To address this gap, we set out to characterize features of the HPI axis and cortisol response in zebrafish larvae following ELC induced by vortex flows. First, we re-examined how ELC in the form of sustained involuntary swimming at 5 dpf affects cortisol response dynamics following a 3-minute vortex at 6 dpf. The rationale for selecting the 5-6 dpf time window is based on a key observation: the machinery for GC synthesis and signalling becomes fully functional around hatching, as evidenced by significant increases in gene expression (Alsop and Vijayan, 2008; Alderman and Bernier, 2009). This corresponds with the gradual rise in whole-body cortisol levels between 2 and 8 dpf. Importantly, cortisol responsiveness to vortex-induced stress peaks at 6 dpf (Castillo-Ramírez et al., 2019), indicating increased developmental sensitivity of the HPI axis at 5–6 dpf. We then examined how ELC impacts the expression and co-expression of *crh*, *avp* (arginine vasopressin), and *oxt* (oxytocin) in the larval NPO. Focusing on these peptides is justified by their established roles in mammals and their conserved functions in stress response and homeostasis in zebrafish. CRH initiates the body’s response to stress and influences autonomic and behavioural responses; AVP plays a crucial role in stress response, social behaviour, and cardiovascular functions, including osmoregulation and blood pressure regulation; and OXT is known for its role in modulating stress and anxiety, social bonding, and reproductive behaviour (Swanson and Sawchenko, 1983; Herman and Cullinan, 1997; Eaton et al., 2008). Assuming that the expression and coexpression profile of NPO cells can undergo stress-derived plasticity, as occurs with PVN cells (Kiss, 1988; Harbuz and Lightman, 1989; Swanson, 1991; Kurrasch et al., 2009), we predicted that ELC may reduce the degree of expression and coexpression of key NPO neuropeptides involved in HPI axis activation.

Next, we explored the effect of ELC at 5 dpf on the transcript levels of genes involved in HPI axis activation, cortisol synthesis and inactivation, and GR signalling, both under baseline conditions and after vortex exposure at 6 dpf. These genes included *hsd11b2* (11-hydroxysteroid dehydrogenase type 2), essential for cortisol inactivation (Krozowski, 1999; Alderman and Vijayan, 2012; Theodoridi et al., 2021); *cyp11c1* (11β-hydroxylase), critical for the synthesis of 11-ketotestosterone and cortisol (Nelson et al., 2013; Tokarz et al., 2015); and *star* (steroidogenic acute regulatory protein), involved in cholesterol transport (Stocco, 2000). We also analysed *crh*, *avp*, *oxt*, *crhr1* (corticotropin-releasing hormone receptor 1), *pomca* (pro-opiomelanocortin, the ACTH precursor), *mc2r* (adrenocorticotropic hormone receptor), *nr3c2* (mineralocorticoid receptor, MR), *nr3c1* (glucocorticoid receptor, GR), and *fkbp5* (FK506 binding protein 5), a stress-responsive regulator of GR (Sinars et al., 2003; Zannas et al., 2016; Hartmann et al., 2021). To explore potential ELC-derived patterns in transcript abundance, we employed a whole-body RNA approach without isolating changes specific to individual HPI axis components (NPO, pituitary, interrenal gland). This exploratory approach is inherently limited, as some target genes are ubiquitously expressed while others are more restricted, with expression often extending beyond the HPI axis.

Despite this limitation, the approach allows us to test whether the overall transcript profile reflects increased cortisol inactivation under both baseline and stress conditions, and whether it indicates an enhanced capacity for cortisol synthesis and stress modulation in larvae previously exposed to ELC. Such patterns would be consistent with an early challenge that imposed prolonged HPI axis activity and elevated cortisol levels.

Lastly, we examined how ELC at 5 dpf affects the cortisol response of 6 dpf larvae to a second vortex applied within the 30-minute refractory period following an initial exposure (Castillo-Ramírez et al., 2024). The refractory period reflects the action of the glucocorticoid feedback loop, during which cortisol levels typically return to baseline and are unresponsive to additional stress. This period has only recently been identified in zebrafish larvae and represents a critical window to assess altered cortisol regulation. In previous work (Castillo-Ramírez et al., 2024), we showed that cortisol levels rise after a 3-minute vortex and return to baseline within 30–40 minutes, depending on the intensity of the vortex. A second vortex administered within 30 minutes of the first fails to elevate cortisol, but blocking GR disrupts this refractory period, indicating GR-dependent feedback. We hypothesized that if ELC at 5 dpf enhances cortisol regulation, larvae exposed to ELC may show an altered response to repeated stress during the refractory period, potentially amplifying the cortisol response to a second vortex. The results are discussed in the context of how ELC influences the initial adjustments of cortisol responsiveness in zebrafish larvae when they encounter environmental challenges for the first time after being in a largely constant environment.

This early-life impact on HPI axis activity further supports the suitability of larval zebrafish as a model for studying the developmental programming of HPA/I axis function.

## RESULTS

### Forced swimming at 5 dpf reduces a larva’s cortisol response to a subsequent exposure to the same type of stress

We first exposed 5 dpf larvae to either vortex flows for 9 hours (ELC larvae) or equal handling without vortex exposure (control larvae) (see Materials and Methods for details). At 6 dpf, we compared the basal whole-body cortisol levels (Fig. 1A) and the cortisol time course following a 3-minute vortex (Fig. 1B) between ELC and control larvae. Consistent with previous data (Castillo-Ramírez et al., 2019), mean baseline whole-body cortisol levels at 6 dpf were similar between control and ELC larvae (Fig. 1C, Mann-Whitney test: U=17, *p*=0.90; N=6 per group), although cortisol levels in ELC larvae were less variable (coefficient of variation: Control, 46.8%, ELC, 18.1%). The 3-minute vortex transiently increased whole-cortisol in both groups, yet ELC larvae showed reduced GC_R_ to the vortex (Fig. 1D, two-way ANOVA on square root-transformed data: group: F(1,50)=35.8, *p*<0.0001; time: F(4,50)=202.9, *p*<0.0001; group x time: F(4,50)=15.4, *p*<0.0001; Bonferroni’s tests: 10’: *p*<0.0001, 20’: *p*<0.0001, 30’: *p*=0.0032, 40’: *p*=0.125, 60’: *p*=0.99; N=6 per group). These results confirm that prolonged forced swimming at 5 dpf reduces a larva’s cortisol response to subsequent homotypic stress, likely due to adjustments in the mechanisms regulating HPI axis reactivity. This raises questions about how these changes relate to the expression and co-expression levels of NPO neuropeptides, which may influence downstream HPI axis activity.

**Figure 1.**
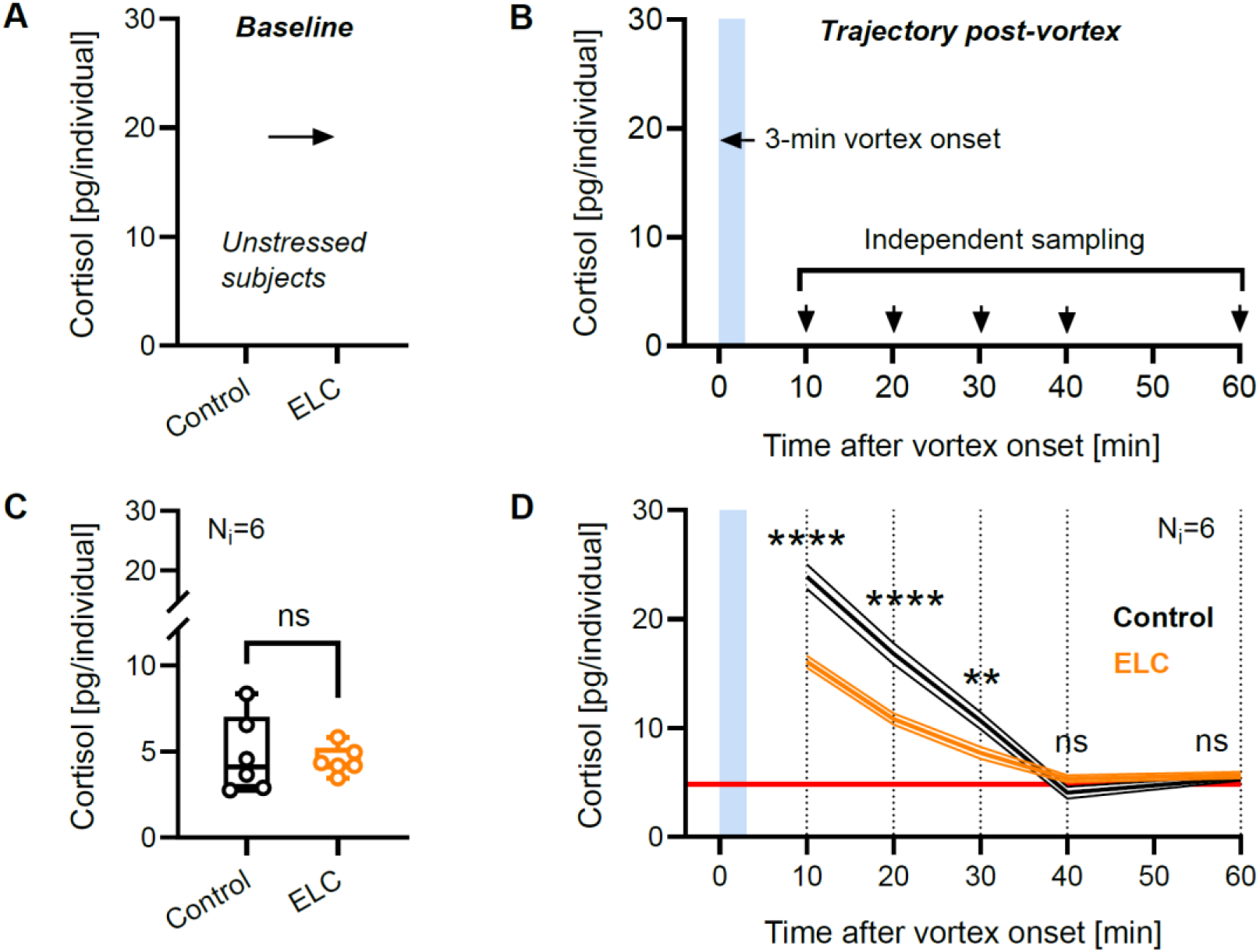
Forced swimming at 5 dpf reduces a larva’s cortisol response to a subsequent exposure to the same type of stress. (**A**,**B**) Comparison steps and time points for measuring baseline whole-body cortisol levels (**A**) and whole-body cortisol trajectory post-vortex (**B**) in 6 dpf larvae with or without ELC exposure. (**C**,**D**) Baseline whole-body cortisol levels (**C**) and whole-body cortisol trajectory post-vortex (**D**) in control (black) and ELC (orange) larvae. (**C**) *p*=0.90 by Mann-Whitney test; N=6 per group. Box and whiskers (min to max), all data points shown. (**D**) 10’ and 20’: \*\*\*\**p*<0.0001, 30’: \*\**p*=0.003, 40’: *p*=0.125, 60’: *p*=0.99 by Bonferroni’s test after a two-way ANOVA on square root-transformed cortisol levels; N=6 per group. Mean ± S.E.M. The red line represents the average baseline whole-body cortisol levels from (**C**).

### NPO imaging shows expression and coexpression of key neuropeptides linked to early life challenge

To examine whether the altered cortisol response to homotypic stress observed in ELC larvae at 6 dpf (Fig. 1) is associated with changes in the expression of *crh*, *avp*, and *oxt* in the larval NPO, we employed multicolor fluorescent *in situ* hybridization at 6 dpf following the ELC protocol implemented at 5 dpf (see Materials and Methods). We compared the numbers of *crh*-, *avp*-, or *oxt*-positive cells in the NPO of unstressed subjects from both groups, ELC and control larvae. Quantification of cell numbers revealed that, compared to control larvae, ELC larvae had fewer *crh*-positive cells within the NPO, as well as fewer *oxt*-positive and fewer *avp*-positive cells (Fig. 2A, Mann-Whitney tests: *oxt*: U=167, *p*=0.03; *avp*: U=112, *p*=0.0004; *crh*: U=85, *p*<0.0001; Group sizes: ELC, N=20, Control, N=27), in line with the idea that ELC can alter the expression of NPO neuropeptides. Additionally, we examined the degree of coexpression between sets of two neuropeptides in the same cell in ELC and control larvae using systematic cell by cell comparisons of pairwise combinatorial ISH staining of *avp*, *crh*, and *oxt* in the larval NPO (Fig. 2B). In both groups, ELC and control larvae, coexpression was consistently high for the combination of *avp* and *crh*, low or moderate for the *crh* and *oxt* combination, and absent or rare for the *oxt* and *avp* combination, in line with previous data (Herget and Ryu, 2015). Also, coexpression was consistently lower in ELC larvae for the combination *avp* and *crh* staining compared to control larvae (Fig. 2C, Mann-Whitney tests: *crh* + *avp*: U=787.5, *p*=0.02; *avp* + *crh*: U=686, *p*=0.002; Group sizes: ELC, N=20, Control, N=27). These results indicate that the changes in GC_R_ observed at 6 dpf in ELC larvae correlate with altered levels of both the expression and coexpression of *avp*, *crh*, and *oxt* in the larval NPO.

**Figure 2.**
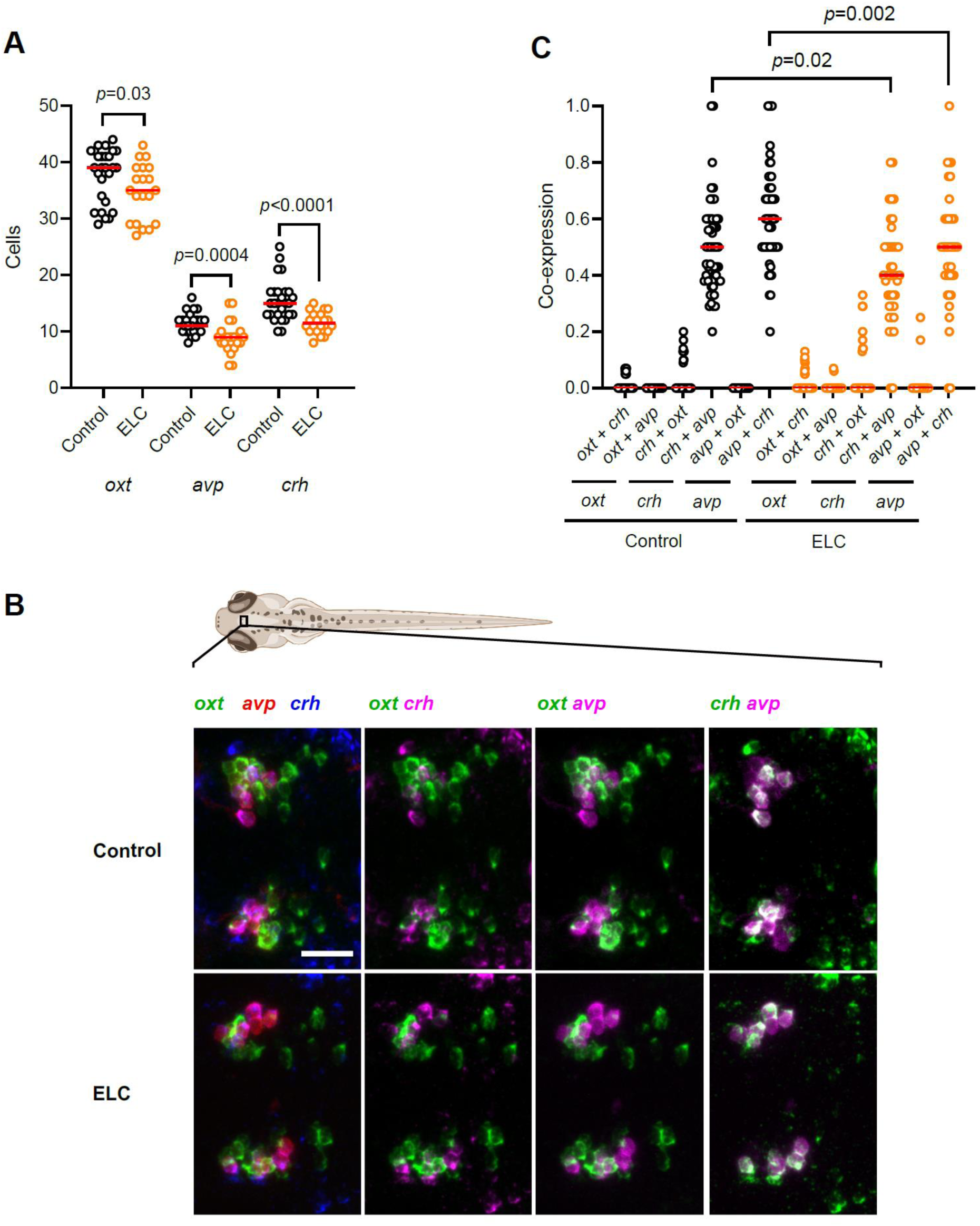
Peptidergic NPO cell numbers in wildtype larvae with or without ELC exposure. (**A**) Comparison of cell numbers in the NPO with or without ELC exposure. (**B**) Maximum intensity projections of confocal stacks show NPO cells expressing *oxt* (green), *avp* (red), or *crh* (blue) in control (above) or ELC (below) larvae. Double-colour comparisons of each pairwise combination are also shown. Scale bar: 25 μm. (**C**) Comparison of coexpression levels in the NPO with or without ELC exposure. (**A**,**C**) Orange: ELC larvae, N=20; black: control larvae, N = 27; all data points shown; *p* values by Mann-Whitney tests. (See also Results.)

### Early life challenge correlates with changes in the transcript abundance of genes involved in cortisol metabolism, steroidogenesis, and stress modulation

Next, we explored whether ELC at 5 dpf is associated with changes in the transcript levels of *hsd11b2*, *cyp11c1*, *star*, *crh*, *avp*, *oxt*, *pomca*, *mc2r*, *crhr1*, *nr3c2*, *nr3c1*, and *fkbp5* at 6 dpf, both at baseline and following a 3-minute vortex. These genes play roles in cortisol synthesis, HPI axis activation, cortisol inactivation, and GR signalling. We employed RT-qPCR to quantify total RNAs from both ELC and control larvae, measuring the fold change in transcript abundance of the 12 genes (Table 1) relative to reference samples from control larvae. Samples were taken at baseline and at 30, 60, and 120 minutes after exposure to the 3-minute vortex (see Materials and Methods). The results indicated that ELC at 5 dpf was associated with altered expression of several genes by 6 dpf (Fig. 3). Compared to controls, ELC larvae showed consistently higher expression of *hsd11b2* under both baseline and stress conditions (Fig. 3A, two-way ANOVA on log-transformed data: group: F(1,32)=53.4, *p*<0.0001; time: F(3,32)=0.06, *p*=0.98; group x time: F(3,32)=0.25, *p*<0.86; Bonferroni’s tests: basal: *p*=0.007, 30’: *p*=0.003, 60’: *p*=0.0006, 120’: *p*=0.013; N=5 per group). Post-stressor, ELC larvae showed elevated expression of *cyp11c1* and *star* (Fig. 3B,C, two-way ANOVAs; *cyp11c1*: group: F(1,32)=20.8, *p*<0.0001; time: F(3,32)=0.99, *p*=0.41; group x time: F(3,32)=1.15, *p*=0.34; Bonferroni’s tests: 60’: *p*=0.007; *star* log-transformed: group: F(1,32)=26.2, *p*<0.0001; time: F(3,32)=2.02, *p*=0.13; group x time: F(3,32)=1.41, *p*=0.26; Bonferroni’s tests: 60’: p=0.014, 120’: p=0.008). Increases in *pomca*, *crh*, and *oxt* expression were also observed post-vortex (Fig. 3D-F; *pomca*: group: F(1,32)=4.99, *p*=0.033; time: F(3,32)=0.45, *p*=0.72; group x time: F(3,32)=2.26, *p*=0.10; Bonferroni’s tests: 120’: *p*=0.038; *crh* log-transformed: group: F(1,32)=6.41, *p*=0.017; time: F(3,32)=1.05, *p*=0.38; group x time: F(3,32)=0.76, *p*=0.52; *oxt* sqrt-transformed: group: F(1,32)=7.24, *p*=0.011; time: F(3,32)=2.65, *p*=0.07; group x time: F(3,32)=1.23, *p*=0.32).

**Figure 3.**
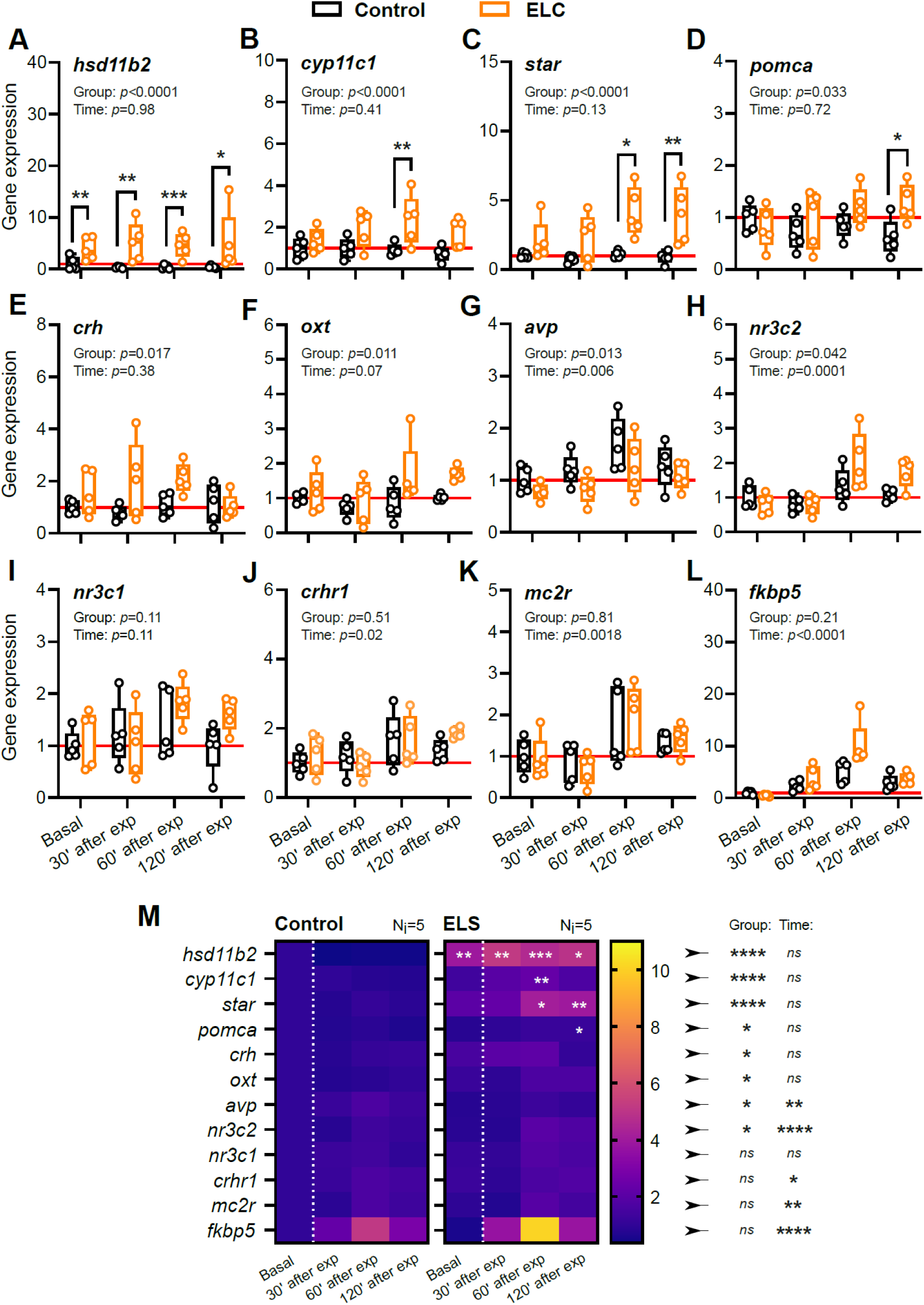
Transcript abundance of genes involved in cortisol metabolism, steroidogenesis, and stress modulation in control and ELC larvae. (**A**-**L**) Fold change in expression of selected genes relative to control reference samples, measured under baseline (basal) conditions and at 30, 60, or 120 minutes after high-strength vortex exposure in control (black) and ELC (orange) larvae; N = 5 per group. *p* values for group and time factors from two-way ANOVAs are shown in the top-left corner of each panel. Asterisks indicate Bonferroni’s test results following the ANOVAs (\**p*<0.05, \*\**p*<0.01, \*\*\**p*<0.001). **(M)** Density grid summarizing fold change comparisons in expression levels of the 12 target genes. Asterisks on the right denote results for the group and time factors from two-way ANOVAs. Asterisks within ELC panels indicate Bonferroni’s test results following the ANOVAs. (See Results for details.)

**Table 1.**
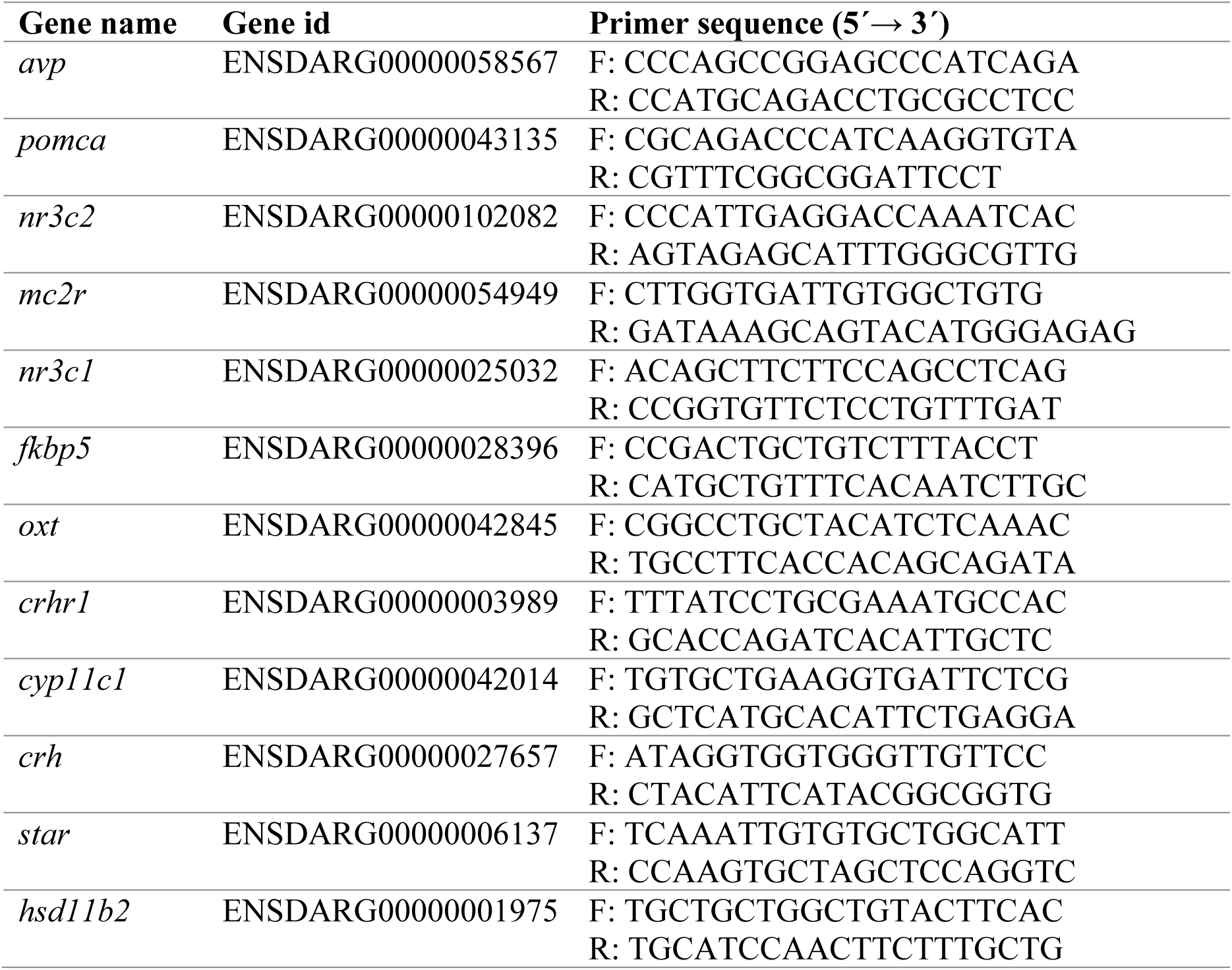
Design primer sequences.

Expression of *avp* and *nr3c2* was influenced by both group and time factors (Fig. 3G,H; *avp*: group: F(1,32)=6.9, *p*=0.013; time: F(3,32)=4.98, *p*=0.006; group x time: F(3,32)=0.23, *p*=0.88; *nr3c2*: group: F(1,32)=4.5, *p*=0.042; time: F(3,32)=9.42, *p*=0.0001; group x time: F(3,32)=2.77, *p*=0.06). Although not statistically significant, *nr3c1* levels tended to increase in ELC larvae post-stressor (Fig. 3I, group: F(1,32)=2.76, *p*=0.11; time: F(3,32)=2.2, *p*=0.11; group x time: F(3,32)=1.2, *p*=0.32). For *crhr1* and *mc2r*, only time influenced expression (Fig. 3J,K; *crhr1*: group: F(1,32)=0.45, *p*=0.51; time: F(3,32)=3.9, *p*=0.02; group x time: F(3,32)=1.1, *p*=0.37; *mc2r*: time: group: F(1,32)=0.06, *p*=0.81; time: F(3,32)=6.2, *p*=0.002; group x time: F(3,32)=0.31, *p*=0.82). Lastly, *fkbp5* transcript levels rose initially and declined later in both ELC and control larvae (Fig. 3L, log-transformed: group: F(1,32)=1.7, *p*=0.21; time: F(3,32)=45.7, *p*<0.0001; group × time: F(3,32)=4.6, *p*=0.009). In summary, we observed differences in transcript levels between control and ELC larvae at 6 dpf. Expression of *hsd11b2* was elevated in both baseline and stress conditions, while *cyp11c1* and *star* increased specifically after vortex exposure. Stressed ELC larvae also showed higher levels of *pomca*, *crh*, and *oxt*. Expression of *avp* and *nr3c2* was influenced by both group and time factors, while trends in *nr3c1*, *crhr1*, and *mc2r* were primarily driven by time following stressor exposure. Finally, *fkbp5* levels initially increased and then declined in both groups (Fig. 3M).

### Early life challenge modifies glucocorticoid reactivity to a second vortex during the refractory period

A reduced capacity of the NPO to influence HPI axis reactivity, along with enhanced cortisol inactivation following ELC, may partly account for the diminished GC_R_ observed after the initial post-ELC vortex. Furthermore, increased cortisol synthesis capacity and the availability of stress modulators could influence GC_R_ dynamics during repeated homotypic stress, potentially affecting the response to a second vortex. To investigate this possibility, we compared whole-body cortisol levels between ELC and control larvae at 6 dpf after two consecutive 3-minute vortices, applied within the 30-minute refractory period following the initial exposure (Castillo-Ramírez et al., 2024) (Fig. 4A). ELC larvae showed increased whole-body cortisol in response to the second vortex applied during this refractory period, whereas control larvae did not (Fig. 4B, Unpaired two-tailed *t*-test: t(10)=13.9, p<0.0001; N=6 per group). Both ELC and control larvae had similar baseline cortisol levels (Fig. 4C, One-sample *t*-test against fold change of ‘1’: t(5)=0.9, p=0.40; N=6 per group; data from Fig. 1C). However, ELC larvae showed a reduced cortisol response to the first vortex as well as an enhanced response to the second vortex (Fig. 4D, two-way ANOVA: group: F(1,20)=9.8, p=0.0052; vortex: F(1,20)=208.1, p<0.0001; group × vortex: F(1,20)=149.5, p<0.0001; Bonferroni’s tests: vortex #1: p<0.0001, vortex #2: p<0.0001; N=6 per group; data from Fig. 1D and Fig. 4B). As a result, control and ELC larvae showed distinct cortisol reactivity to the second vortex. While ELC larvae showed a GC_R_ to the second vortex that was 91% of their response to the first vortex, control larvae showed only 18% (Fig. 4E, Unpaired two-tailed *t*-test; t(10)=20.7, p<0.0001; N=6 per group; data from Fig. 4D). These findings indicate that prolonged forced swimming at 5 dpf improves the ability of larvae at 6 dpf to regulate cortisol levels and adjust HPI axis activity in response to stress, enhancing their capacity to cope with repeated exposure to the same stressor.

**Figure 4.**
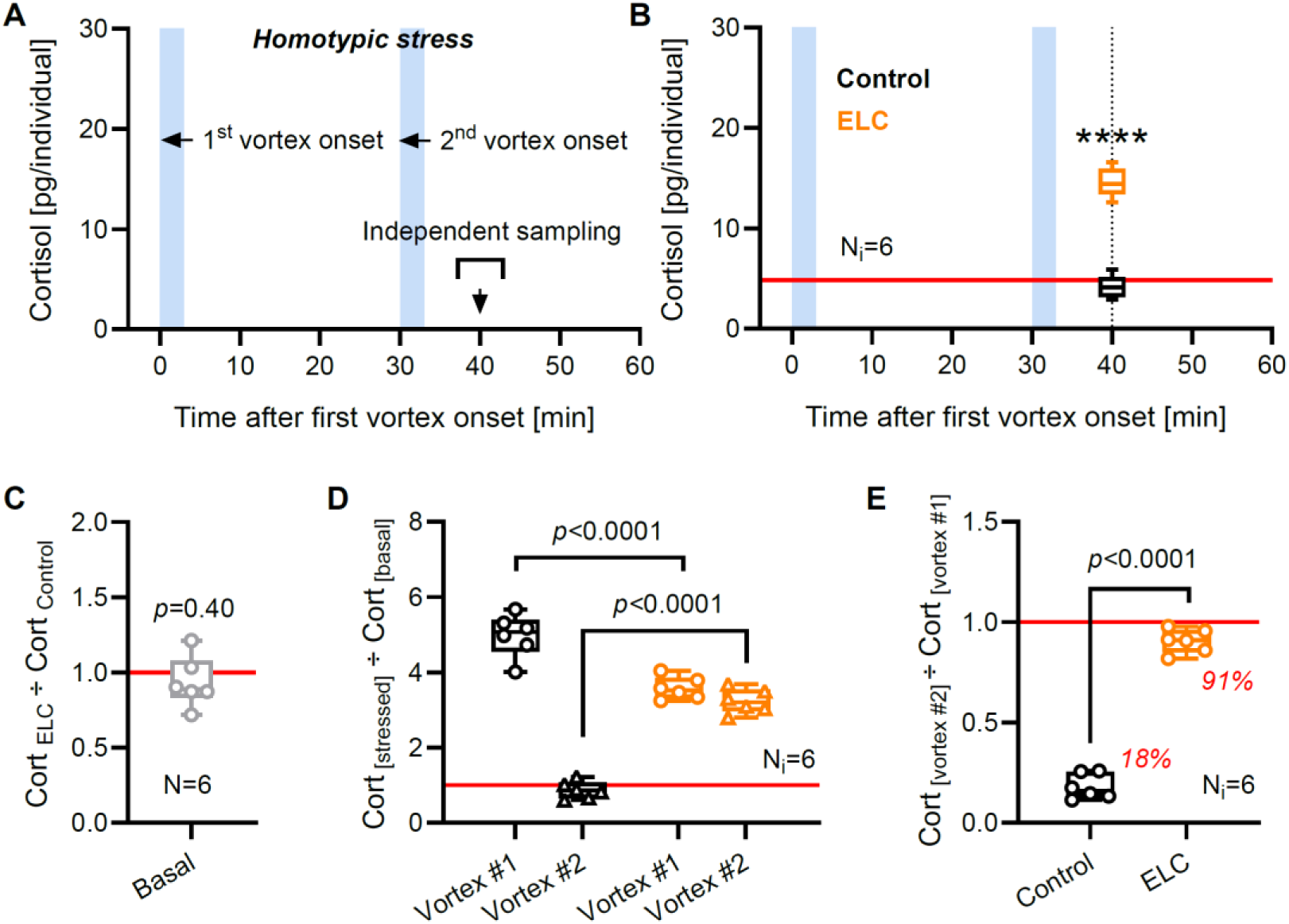
ELC alters GC_R_ to a second vortex during the refractory period. (**A**) Sampling scheme for measuring GC_R_ to homotypic stress in 6 dpf larvae with or without ELC. (**B**) Whole-body cortisol following two 3-minute vortices applied within a 30-minute refractory period in control (black) and ELC (orange) larvae. Control larvae did not show cortisol levels above baseline in response to a second 3-minute vortex, while ELC larvae did. N=6 per group. \*\*\*\**p* < 0.0001 by unpaired two-tailed *t*-test. (**C**) Baseline whole-body cortisol levels in ELC larvae relative to controls (data from Fig. 1C), depicted for the sake of interpreting panels (**D**) and (**E**). *p* = 0.40 by one-sample *t*-test against fold change of ‘1’. (**D**) GC_R_ to 1^st^ and 2^nd^ vortices relative to baseline in control and ELC larvae. Data from Fig. 1C, Fig. 1D (10 min post-vortex), and (**B**). N=6 per group. *p* values indicate Bonferroni’s test results following a two-way ANOVA. (**E**) GC_R_ to the 2^nd^ vortex relative to GC_R_ to the 1^st^ vortex in control (black) and ELC (orange) larvae (data from **D**). *p* < 0.0001 by unpaired two-tailed *t*-test. (**B**-**E**) Box and whiskers (min to max), (**C-E**) all data points shown.

## DISCUSSION

Here we provide strong evidence for extended effects of early forced swimming on cortisol regulation and stress responsiveness in zebrafish larvae. We observed that ELC at 5 dpf resulted in significant alterations in cortisol reactivity to subsequent stress events. Specifically, ELC larvae had similar baseline cortisol levels compared to controls but showed reduced GC_R_ to a single vortex at 6 dpf. This finding is consistent with previous data (Castillo-Ramírez et al., 2019) and reveals adaptive adjustments in stress axis function following prolonged early-life stressor exposure. Notably, while ELC larvae showed reduced GC_R_ to the initial stressor, they had increased cortisol responsiveness to a second stress event occurring within the 30-minute refractory period following the first vortex (Castillo-Ramírez et al., 2024). This dual response pattern suggests that ELC larvae have enhanced stress responsiveness during homotypic stress, potentially due to adaptive changes in HPI axis signalling and cortisol metabolism.

Our findings are consistent with research in fish and other vertebrates showing that early life stress and GC manipulation can have lasting effects on HPA/I axis function and stress responsiveness. In larval zebrafish, optogenetic techniques enable non-invasive control of endogenous GC levels (De Marco et al., 2013; 2016). Transgenic larvae expressing Beggiatoa photoactivated adenylyl cyclase (bPAC) (Ryu et al., 2010; Stierl et al., 2011) under a StAR promoter can induce light-dependent activation of interrenal gland steroidogenic cells, resulting in elevated cortisol levels (Gutierrez-Triana et al., 2015).

Persistent activation of these cells during early development leads to chronically high cortisol levels and a diminished cortisol response to later stressors (Nagpal et al., 2024). Additionally, optogenetically elevated GC levels early in life disrupt hypothalamic neurogenesis, causing precocious development, failed maturation, and impaired feeding and growth (Eachus et al., 2024). Chronic stress induced by random shocks in developing zebrafish also causes sustained high cortisol levels and increased expression of glucocorticoid and mineralocorticoid receptors, which correlate with increased anxiety-like behaviour and elevated cortisol levels later in life (Chin et al., 2022). Zebrafish exposed to air-exposure stress at different developmental stages showed increased cortisol levels, altered stress responses, and changes in ion concentrations (Hare et al., 2021). Similar trends are observed in other species. In rainbow trout, chronic cortisol intake and daily stress exposure reduce GC_R_ (Barton et al., 1987). Rodent studies show that neonatal rats treated with dexamethasone show reduced GC_R_ in response to cold and restraint stress later in life (Felszeghy et al., 2000), while mice subjected to prolonged low-dose corticosterone exposure show suppressed GC_R_, adrenal atrophy, and decreased CRH mRNA levels in the PVN (Kinlein et al., 2015). In humans, infants exposed to high cortisol levels *in utero* show elevated baseline cortisol and reduced GC_R_ in response to separation stress (O’Connor et al., 2013). Collectively, these observations underscore the complex interplay between ELC, GC regulation, and stress responsiveness early in life. However, the mechanisms through which ELC affects HPA/I axis function and leads to long-term changes in stress response patterns are still largely unclear, particularly regarding the initial phase of this interaction.

The HPA axis regulates its stress-induced activation through negative feedback mechanisms (Dallman and Yates, 1969; Dallman et al., 1994; Dallman, 2005; Shipston, 2022). In mammals, this feedback acts on the PVN, where GCs suppress the expression of stress modulators like *crh* (Malkoski et al., 1999). Given the structural and chemoarchitectural similarities between the mammalian PVN and the zebrafish NPO (Herget et al., 2014), it is likely that the NPO is subject to similar GC-mediated regulatory feedback (Castillo-Ramírez et al., 2024). Supporting this, zebrafish with disrupted GC signalling show increased numbers of *crh*-expressing cells in the NPO (Ziv et al., 2013; Facchinello et al., 2017). In mammals, regulation of the PVN involves not only the expression of individual modulators like *crh* but also the co-expression of multiple neuropeptides within the same cells (Sawchenko et al., 1984; Whitnall et al., 1985; Simmons and Swanson, 2009; Biag et al., 2012). However, in the zebrafish NPO, neuropeptides tend to be produced by distinct, tightly packed cells, with limited co-expression, although certain combinations show occasional to moderate co-expression (Herget and Ryu, 2015). Given that stress can induce plastic changes in the expression and co-expression of neuropeptides in the PVN (Kiss, 1988; Harbuz and Lightman, 1989; Swanson, 1991; Kurrasch et al., 2009), we hypothesized that ELC might similarly alter the expression and co-expression profiles of key neuropeptides in the NPO. Our findings show that ELC larvae had fewer *crh*-, *avp*-, and *oxt*-positive NPO cells than controls, with a marked reduction in *crh* and *avp* co-expression. These results suggest that, analogous to the mammalian PVN (Ziv et al., 2013; Herget et al., 2014; Facchinello et al., 2017), the zebrafish NPO serves as a critical site for GC feedback regulation. Moreover, they indicate that ELC induces plasticity in neuropeptide expression within the developing NPO, potentially affecting downstream HPI axis activity. The reduced number of *crh*- and *avp*-positive cells in the NPO may contribute to the reduced GC_R_ observed following the initial stress exposure in ELC larvae.

While the expression profile of steroidogenic enzymes involved in GC biosynthesis has been characterized in developing zebrafish (Weger et al., 2018), our data revealed group differences between ELC and control larvae. Specifically, ELC larvae showed upregulation of *hsd11b2* under both baseline and stress conditions, as well as increased expression of *cyp11c1*, *star*, *pomca*, *crh*, *oxt*, *avp*, and *nr3c2* following a 3-minute vortex. Additionally, transcripts for *avp*, *nr3c2*, *crhr1*, *mc2r*, and *fkbp5* showed time-dependent changes post-stressor. Interestingly, previous studies in fish have linked whole-body cortisol levels with *hsd11b2* and *cyp11c1* mRNA expression (Tsalafouta et al., 2014), while prolonged cortisol release has been observed in zebrafish lacking functional Hsd11b2 (Theodoridi et al., 2021). Notably, although ELC larvae showed fewer *crh*-positive cells in the NPO, their whole-body *crh* levels increased after acute stress. This suggests that ELC may influence the maturation of stress-related circuits involving NPO *crh*-positive cells (Bolton et al., 2022). While whole-body qPCR cannot isolate transcript changes specific to the HPI axis, given that some genes are ubiquitously expressed while others have restricted but broader distribution, the overall transcript pattern in 6 dpf ELC larvae points to enhanced cortisol inactivation under both baseline and stress conditions. Additionally, the upregulation of steroidogenic enzymes and stress modulators indicates an increased capacity for cortisol synthesis and regulation during stress. These transcriptional changes, alongside altered *avp*, *crh*, and *oxt* expression and co-expression patterns in the NPO, suggest that ELC enhances cortisol regulation, preventing excessive cortisol release and promoting an appropriate response to homotypic stress (Fig. 5).

**Figure 5.**
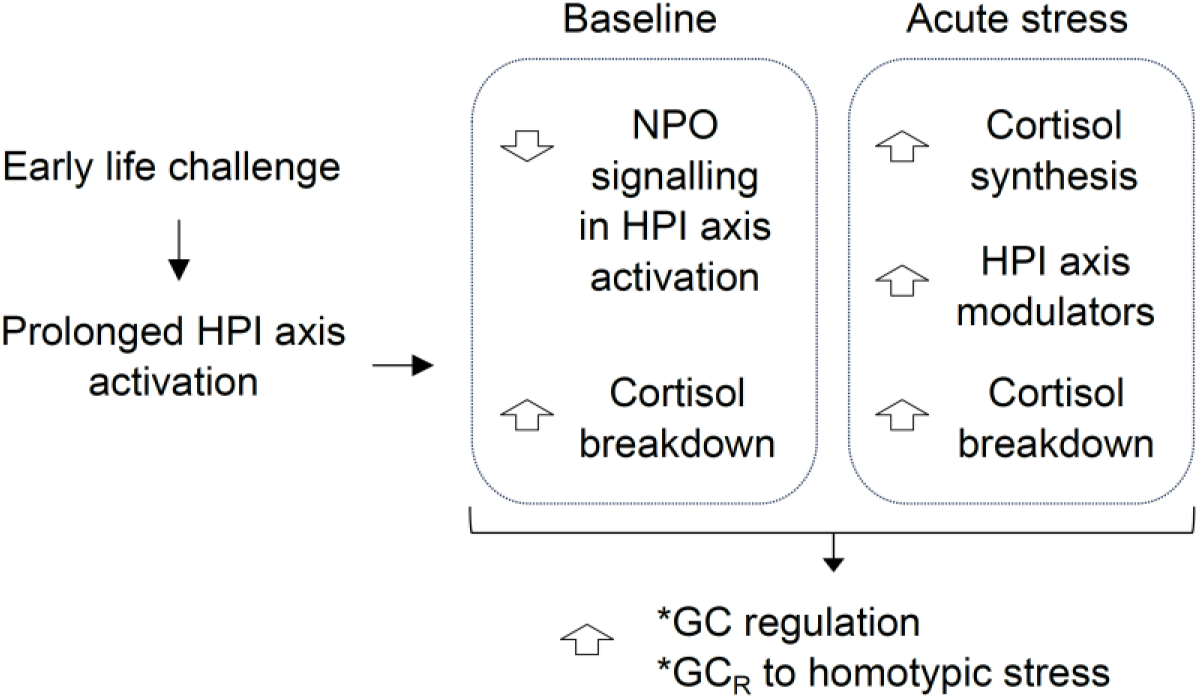
Proposed model illustrating the initial adaptation of the HPI axis following ELC. Prolonged early-life activation of the HPI axis, induced by increased environmental challenge, diminishes the NPO’s capacity to influence downstream HPI axis activity. Baseline conditions show enhanced cortisol inactivation, which can persist upon stress onset. There is an increase in cortisol synthesis capacity and elevated levels of stress modulators. These adaptations enhance the larva’s capability to regulate GC levels, preventing excess cortisol while facilitating its production in response to familiar stress.

Our findings contribute to a better understanding of how early-life environments shape GC_R_, NPO plasticity, and GC pathways during the early stages of HPI axis development. They also highlight the value of larval zebrafish as a model organism for studying the developmental programming of HPI axis function (Weinstock, 2008; Glover et al., 2010; Moisiadis and Matthews, 2014; de Abreu et al., 2021; Eachus et al., 2021; Swaminathan et al., 2023). While this study focused primarily on cortisol dynamics, future research should compare forced swimming in vortex conditions to a larva’s routine swimming patterns, investigating the broader physiological effects of vortex-induced stress, which offers an excellent tool for high-throughput testing. Additional studies could evaluate measures such as whole-body lactate and cholesterol levels, cardiac activity, and anaerobic metabolism to provide a more comprehensive understanding of the physiological demands imposed by vortex exposure. Moreover, examining how prolonged early-life forced swimming affects cortisol production in interrenal cells (Wilson et al., 2016), the maturation of stress-related circuits in the NPO, and stress responses in later life stages could reveal long-lasting impacts on cortisol regulation and stress-related pathways. Such research would clarify how vortex exposure as an early-life challenge influences not only immediate stress responses but also long-term physiological and behavioural outcomes, offering valuable insights into stress-related developmental programming.

## MATERIALS AND METHODS

### Zebrafish husbandry, handling, and experimental unit

Zebrafish breeding and maintenance were conducted under standard conditions (Westerfield, 2000). Groups of thirty wild-type eggs (cross of AB and TL strains, AB/TL) were collected in the morning, and the embryos were raised on a 12:12 light/dark cycle at 28°C in 35 mm Petri dishes with 5 ml of Embryo-medium2 (E2). The E2 medium (0.5x E2, 1 L) consisted of 5 mM NaCl, 0.25 mM KCl, 0.5 mM MgSO_4_ × 7 H_2_O, 0.15 mM KH_2_PO_4_, 0.05 mM Na_2_HPO_4_, 0.5 mM CaCl_2_, and 0.71 mM NaHCO_3_. At 3 dpf, the E2 medium was renewed, and chorions and debris were removed from the dishes. Experiments were performed with 5-6 dpf larvae. In all experiments, each experimental unit (replicate) comprised a group of thirty larvae, maintained in a 35 mm Petri dish. All dishes were kept under identical conditions in the incubator on top of the stirrer plate (see below) to ensure no perturbation. Zebrafish experimental procedures were conducted according to the guidelines of the German animal welfare law and were approved by the local government (Regierungspräsidium Karlsruhe; G-29/12).

### Water vortex flows

We used water vortices in a high-throughput manner to induce rheotaxis and cortisol elevation (Castillo-Ramírez et al., 2019). Groups of thirty larvae (either 5 or 6 dpf, depending on the experiment) were placed in 35 mm Petri dishes containing 5 ml of E2 medium (experimental units) and exposed to controlled vortices generated by the spinning movements of small encapsulated (coated) magnetic stir bars (6 x 3 mm, Fisherbrand, #11888882, Fisher Scientific, Leicestershire, UK) inside the dishes. Each Petri dish, either containing a stir bar or not (depending on the experimental group), was always positioned on a magnetic stirrer plate (Variomag, Poly 15; Thermo Fisher Scientific, Leicestershire, UK) and maintained at 28°C in an incubator (RuMed 3101, Rubarth Apparate GmbH, Laatzen, Germany). Vortex flows were induced by the movements of the stir bars, driven by magnetic field inversions of the stirrer plates set at 330 revolutions per minute (rpm). The magnetic field inversions alone did not affect whole-body cortisol levels in larval zebrafish, and we limited the speed to 330 rpm to avoid potential ceiling effects in vortex-dependent cortisol increases, as previously reported (Castillo-Ramírez et al., 2019). At 5 dpf, larvae were either exposed to 9 hours of continuous vortices (ELC larvae) or to magnetic field inversions without vortices (Control larvae), as no stir bars were present in the control group’s Petri dishes. The 9-hour exposure period was designed to maximize the duration of vortex exposure while adhering to the light/dark cycle. Larvae at 5 dpf exposed to vortex flows for 9 hours show elevated whole-body cortisol levels, peaking shortly after vortex initiation and remaining high for up to four hours compared to controls, i.e., unexposed larvae that are handled similarly but without vortex flows in their medium. By six hours after vortex onset, cortisol levels in both exposed and control larvae are comparable.

Additionally, exposed larvae consistently engage in positive rheotaxis throughout the vortex exposure, as indicated by their body angle relative to the incoming current, even 8.5 hours after the vortex began (Castillo-Ramírez et al., 2019). At 6 dpf, larvae previously exposed (ELC) or not exposed (Control) to the 9-hour forced swimming period at 5 dpf were used for the following analyses: 1) measurement of baseline whole-body cortisol levels (Fig. 1C), extraction of total RNA for quantification of baseline transcript abundance of 12 genes, or whole-mount fluorescent *in situ* hybridization and immunohistochemistry to assess cell number and coexpression (Fig. 2); 2) measurement of whole-body cortisol at 10, 20, 30, 40, and 60 minutes after a single 3-minute vortex (330 rpm) (Fig. 1D), or total RNA extraction to quantify transcript abundance of the 12 genes post-vortex (Fig. 3); and 3) measurement of whole-body cortisol after a second 3-minute vortex administered 30 minutes after the first (Fig. 4). The selected 10, 20, 30, 40, and 60-minute intervals (Fig. 1D) effectively captured cortisol dynamics following the initial stressor, providing a broad range to account for variability in stress response timing. This design ensured experimental rigor while adhering to ethical guidelines for subject use. At 6 dpf, larvae were immobilized in ice water before cortisol detection or total RNA extraction (see below).

### Whole-body cortisol

At 6 dpf, groups of 30 ELC or Control larvae (replicates), either unexposed to the 3-minute vortex (baseline measurement) or exposed to one or two vortices applied within 30 minutes, were immobilized in ice water (<2°C) to minimize stress. Excess water was removed, and larvae were immediately frozen in an ethanol/dry-ice bath to ensure rapid euthanasia and preservation. Samples were then stored at −20°C for subsequent cortisol extraction, which took place between 10:30 and 11:30 hours. The procedures for cortisol measurements and the homemade ELISA were as previously described (Yeh et al., 2013). The homemade ELISA used for cortisol detection was validated through assessments of intra-assay and inter-assay precision, recovery rates, and cross-reactivity. Results across batches and sessions were consistent, with no significant differences in baseline cortisol levels, further confirming the reliability of the assay. Full details of the validation process, including comparisons with a commercial kit, can be found in Castillo-Ramírez et al. (2024).

### Quantitative Real-Time PCR (qPCR)

Total RNA was extracted from groups of thirty larvae, as previously described for cortisol extraction. RNA isolation was performed using the RNeasy Micro Kit (Qiagen, Hilden, Germany), according to the manufacturer’s protocol. The purity and concentration of RNA were determined using a NanoDrop spectrophotometer (Thermo Fisher Scientific), and the integrity of the RNA was confirmed using an RNA 6000 Pico Kit chip (Agilent Technologies) on the Agilent 2100 Bioanalyzer. Only samples with RNA Integrity Numbers (RIN) above 7.0 were used in subsequent qPCR analysis. qPCR reactions were carried out using the Power SYBR Green RNA-to-Ct 1-Step Kit (Thermo Fisher Scientific, Leicestershire, UK) on a 7500 Real-Time PCR system (Applied Biosystems). RNA was reverse transcribed and amplified in a single-step reaction. Primers for the target genes (listed in Table 1) were either designed in-house using Primer3 software or obtained from published sources. Primer efficiency was confirmed using a standard curve approach, ensuring that only primers with an efficiency of 90% to 110% were used in further experiments. The gene Elongation Factor 1-alpha (EF1α) (F: CTGGAGGCCAGCTCAAACGT R: ATCAAGAAGAGTAGTACCGCTA) was selected as the reference gene based on its known stability in zebrafish across various tissues and treatments (McCurley and Callard, 2008). To ensure that EF1α maintained stable expression in our experimental setup, the cycle threshold (Ct) values of EF1α were examined across all experimental groups. Consistency in Ct values across conditions indicated stable expression of EF1α, validating its use as a reliable internal control for normalizing the expression of target genes. Minimal variation in Ct values across all treatments and replicates demonstrated that the gene was unaffected by experimental conditions. Each 20 µL qPCR reaction contained 10 µL of SYBR Green Master Mix, 0.5 µL of forward and reverse primers (10 µM), 0.4 µL of ROX reference dye, and 100 ng of total RNA. The thermal cycling program included an initial reverse transcription step at 48°C for 30 min, followed by denaturation at 95°C for 10 min, and 40 cycles of 95°C for 15 s and 60°C for 1 min.

Relative gene expression levels were calculated using the 2^−ΔΔCt method (Livak and Schmittgen, 2001), with EF1α serving as the reference gene for normalization. The fold change in gene expression was calculated by comparing the treated samples to the control group, and statistical analyses were performed on the ΔCt values to determine significance.

### Whole-mount fluorescent *in situ* hybridization, immunohistochemistry, and imaging

Whole-mount fluorescent *in situ* hybridization and immunohistochemistry were performed as described elsewhere (Kastenhuber et al., 2010; Lauter et al. 2011), using riboprobes for *avp* (Eaton et al., 2008), *oxt* (Unger and Glasgow, 2003), and *crh* (Löhr et al., 2009), and a primary chicken antibody labeling GFP (1:500, Abcam), with a secondary anti-chicken Alexa 488 antibody (1:1000, invitrogen). For imaging, specimens were cleared in 80% glycerol (Gerbu Biotechnik GmbH, Heidelberg, Germany) in PBS for 1 h. Confocal stacks were recorded using a Leica SP5 confocal microscope (Leica Microsystems GmbH, Wetzlar, Germany) with a Nikon 20x glycerol objective (Nikon, Tokyo, Japan). Each channel was recorded sequentially to reduce interfering signals from overlapping emission spectra. Zoom, dimensions, gain, offset, average, and speed were adjusted for each stack to obtain the optimal image quality of the desired volume. Stacks were evaluated using Amira 5.4 (Thermo Fisher Scientific, Leicestershire, UK) to create maximum intensity projections, which were spatially restricted to the volume of interest, excluding signals from planes in front or behind. Brightness and contrast were adjusted for each channel.

### General design and statistical analysis

Cortisol and RT-qPCR measurements were conducted on distinct groups of thirty larvae (replicates), with larval density per well kept constant. For each measurement, all thirty larvae in a well were used. Each replicate was fully independent of the others. For cell counting measurements, each replicate consisted of a single larva. In all experiments, treatments were randomly assigned to replicates, and blinding was implemented. An initial experimenter conducted the treatments, collected, and labelled the samples. A second experimenter then performed the measurements on the labelled samples, assigning new labels. The first experimenter subsequently quantified the results using these newly encoded samples. Our sample sizes are consistent with those commonly used in the field and align with previous publications (Castillo-Ramírez et al., 2019; De Marco et al., 2013; Yeh et al., 2013; De Marco et al., 2016; vom Berg-Maurer et al., 2016; Herget et al., 2023;). They are based on prior work that established acceptable coefficients of variation for measurements while minimizing the use of subjects, in accordance with ethical guidelines. Data were tested for normality and homoscedasticity using the Shapiro-Wilk and KS normality tests and the Brown–Forsythe and Bartlett’s tests, respectively. We employed Mann-Whitney tests for pairwise comparisons involving non-normally distributed data, two-way ANOVAs followed by Bonferroni’s post-hoc tests for multiple comparisons. In cases where the data did not meet the assumption of homoscedasticity, we applied log transformations to normalize variance. Unpaired two-tailed t-tests were used for comparing independent groups, and one-sample t-tests were performed when comparing data to a hypothetical mean (i.e., a fold change of 1). No data points or samples were excluded from the analysis. Statistical analyses were performed using MS-Excel (Microsoft Corp; Redmond, WA, USA) and Prism 10.2.0 (Graphpad Software Inc, San Diego, CA, USA).

## Acknowledgments

We would like to thank L. Flores-García for assistance with the experiments, Karl J. Iremonger and Chrysanthi Fergani for providing useful comments on a previous draft, R. Singer and A. Schoell for expert fish care, and R. Rödel for technical support.

## Author Contributions

L.A.C-R., Conceptualization, Data curation, Formal analysis, Investigation, Validation, Methodology, Writing - original draft; U.H., Data curation, Formal analysis, Methodology, Writing – original draft; S.R., Conceptualization, Resources, Supervision, Funding acquisition, Methodology, Project administration, Writing – review; R.J.DM., Conceptualization, Resources, Supervision, Funding acquisition, Investigation, Validation, Methodology, Data curation, Formal analysis, Visualization, Project administration, Writing - original draft, Writing - review and editing.

## Data availability

The experimental datasets presented in this study are available upon request from the authors.

